# Gaze-evoked deformations of the optic nerve head in thyroid eye disease

**DOI:** 10.1101/2020.03.19.999706

**Authors:** Liam K. Fisher, Xiaofei Wang, Tin A. Tun, Hsi-Wei Chung, Dan Milea, Michaël J.A. Girard

## Abstract

**Purpose:** To assess gaze evoked deformations of the optic nerve head (ONH) in thyroid eye disease (TED), using computational modelling and optical coherence tomography (OCT).

**Methods:** Multiple finite element models were constructed: One model of a healthy eye, and two models mimicking effects of TED; one with proptosis and another with extraocular tissue stiffening. Two additional hypothetical models had extraocular tissue softening or no extraocular tissue at all. Horizontal eye movements were simulated in these models.

OCT images of the ONH of 10 healthy volunteers and 1 patient with TED were taken in primary gaze. Additional images were recorded in the same subjects performing eye movements in adduction and abduction.

The resulting ONH deformation in the models and human subjects was measured by recording the ‘tilt angle’ (relative antero-posterior deformation of the Bruch’s membrane opening). Effective stress was measured in the peripapillary sclera of the models.

**Results:** In our computational models the eyes with proptosis and stiffer extraocular tissue had greater gaze-evoked deformations than the healthy eye model, while the models with softer or no extraocular tissue had lesser deformations, in both adduction and abduction. Scleral stress correlated with the tilt angle measurements.

In healthy subjects, the mean tilt angle was 1.46° ± 0.25 in adduction and −0.42° ± 0.12 in abduction. The tilt angle measured in the subject with TED was 5.37° in adduction and −2.21° in abduction.

**Conclusions:** Computational modelling and experimental observation suggest that TED can cause increased gaze-evoked deformations of the ONH.

## Introduction

Thyroid eye disease (TED) is an autoimmune disorder where activated orbital fibroblasts differentiate into adipocytes or myofibroblasts and have increased hyaluronan production (1). These processes often involve expansion and infiltration of the orbital fat and the extraocular muscles (2, 3). In addition to an aesthetic burden (eyelid retraction, proptosis), the mechanical disturbance of the peri-ocular tissues can cause serious ophthalmic dysfunction (diplopia, dry eyes, elevated intra-ocular pressure) (4). Sight threatening dysthyroid optic neuropathy (DON) can also occur in severe TED. This neuropathy is typically attributed to extraocular muscle expansion compressing the optic nerve, especially at the narrow apex of the orbit where there is limited space to accommodate the volume of inflamed tissue (5).

Additionally, an association has been observed between TED and glaucoma. In some cases, this can be attributed to elevated intraocular pressure (IOP) caused by TED (6). However, there also appear to be several TED cases where patients develop neuropathic glaucoma-like symptoms despite having normal IOP and no radiological evidence of compressed optic nerves (7, 8). There is currently no compelling explanation for this finding.

Recent research has proposed that eye movements may have a mechanical effect on the tissues of the optic nerve head (ONH). This theory has been supported by both computational (9-11) and experimental (12-22) findings. Comparison between eye movement and high-IOP conditions suggests that eye movements could contribute to the development of glaucoma, as the stresses and strains induced by eye movement at the ONH (so-called ‘gaze-evoked deformations’) are similar to or greater than those induced by harmful IOP elevations.

The influence that TED may have on these gaze-evoked ONH deformations has not been previously explored. It is plausible that TED affects these deformations, for example proptosis may increase the traction force at the ONH by stretching the optic nerve in the axial direction. The aim of this study is to investigate whether TED exacerbates gaze-evoked ONH deformations using in-vivo and computational techniques.

## Methods

In this study, we used finite element (FE) modelling to better understand the mechanical insult at the level of the ONH during eye movements in TED. Simulated horizontal eye movements (adductions and abductions^1^) were applied to FE models of the orbit, and the resulting mechanical effects (deformation and stress) were assessed in the ONH tissues. We attempted to verify these findings by comparing them to in vivo ONH deformation data from a group of healthy subjects and a TED patient.

### Finite-Element Geometry of the Ocular and Orbital Tissues

Our 3D eye models were adapted and modified from a previous study (**Figure 1**) (9). The optic nerve and eye globe were reconstructed from magnetic resonance imaging (MRI) of a healthy subject. The corneo-scleral shell was assumed to be spherical (outer diameter: 24 mm; thickness: 1 mm) and the optic nerve consisted of three layers: nerve tissue (diameter 2.3mm), the pia mater (thickness: 0.06 mm), and the dura mater (thickness: 0.3 mm). An ONH geometry was constructed at the intersection of the nerve and the corneo-scleral shell, including the scleral flange (length: 0.4 mm; thickness: 0.45 mm), the LC (central thickness: 0.28 mm; anterior and posterior radii of 0.89 and 1.01 mm, respectively), and the prelaminar tissues (thickness: 0.2mm).

**Fig 1:**
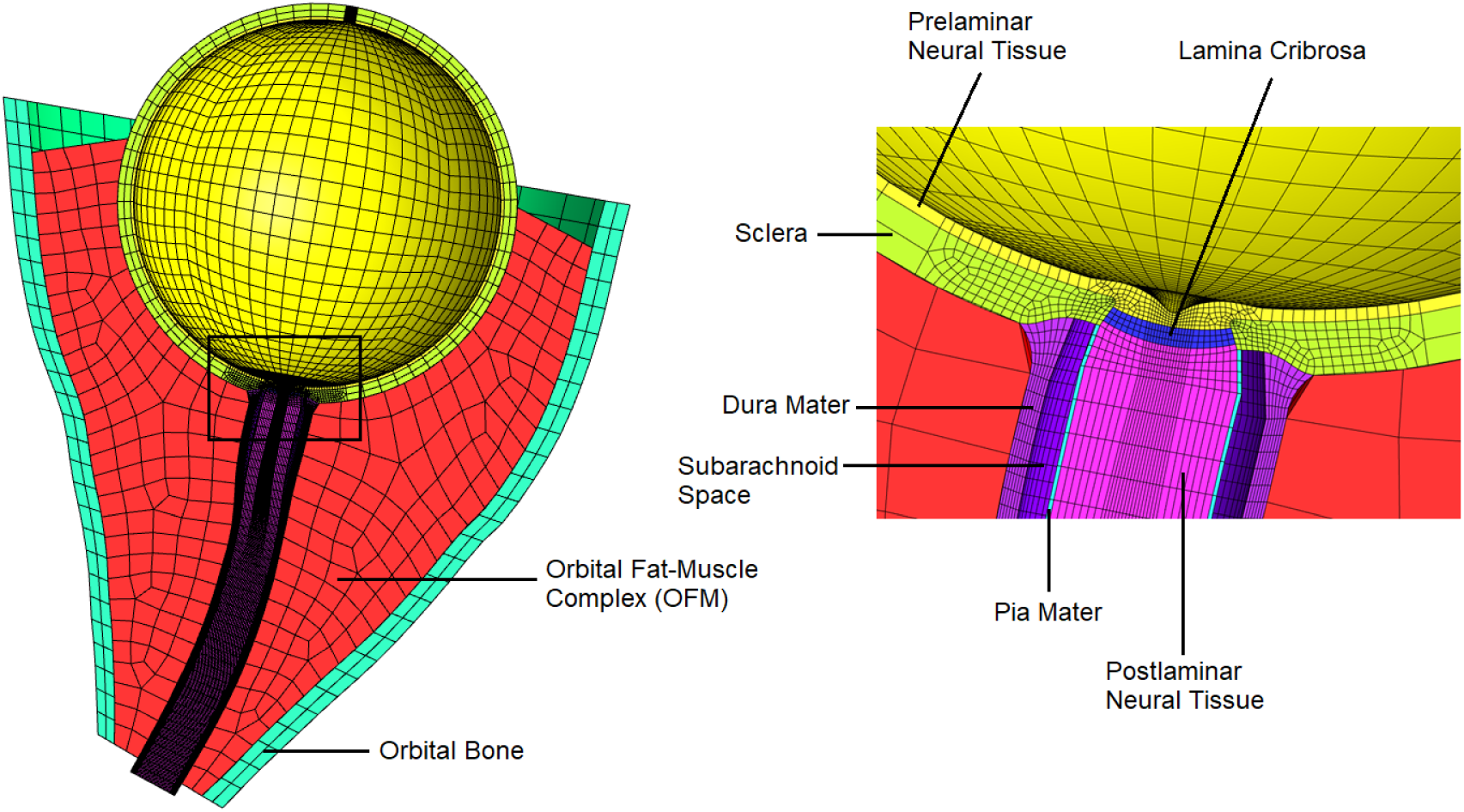
Geometry and mesh of the finite element models (baseline gaze position). The magnified view provides a detailed illustration of the ONH region.

These ocular tissues were enclosed within a boundary structure representing the orbital bone, with an anterior opening of 35.2 mm diameter, an apical opening of 6.1 mm diameter, and depth of 38.3 mm (measured from the midpoint of the medial and lateral orbital rim to the midpoint of the medial and lateral rim of the optic canal). The space between this boundary and the ocular and nerve structures was filled with a material described as the orbital fat-muscle complex (OFM). For simplicity, extraocular muscles were not represented by discrete structures. Our analysis was focused on the mechanical environment of the ONH and the immediate orbital fat, and eye movements were induced artificially, so individual muscle bodies were not considered to be necessary.

This geometry was discretized into a mesh with 55084 eight-node hexahedra using ICEM CFD (ANSYS, Inc., Canonsburg, PA, USA). The mesh density was numerically validated with a convergence test. The eye and orbit were assumed to be symmetrical about the transverse plane, allowing the entire orbit to be described by a half-model with appropriate boundary conditions.

### Material Properties of the Ocular and Orbital Tissues

The properties for the ocular tissues in our models were largely adapted from our previous work. The sclera (23) and lamina cribrosa (24) were modelled as fibre-reinforced composites. Neural tissue and the orbital bone were modelled as isotropic elastic materials and thus characterised by a single stiffness value. The stiffness value for neural tissue was obtained from the literature (25) and the stiffness value of bone was set at an arbitrarily high value to account for its relative rigidity compared to the soft tissues. The pia and dura were modelled as Yeoh materials, derived from experimental data from porcine eyes (9). These parameters are listed in **Table 1**. While the above material properties were common to all models, the properties of the OFM varied. These differences are explained in a later section of this paper.

**Table 1:**
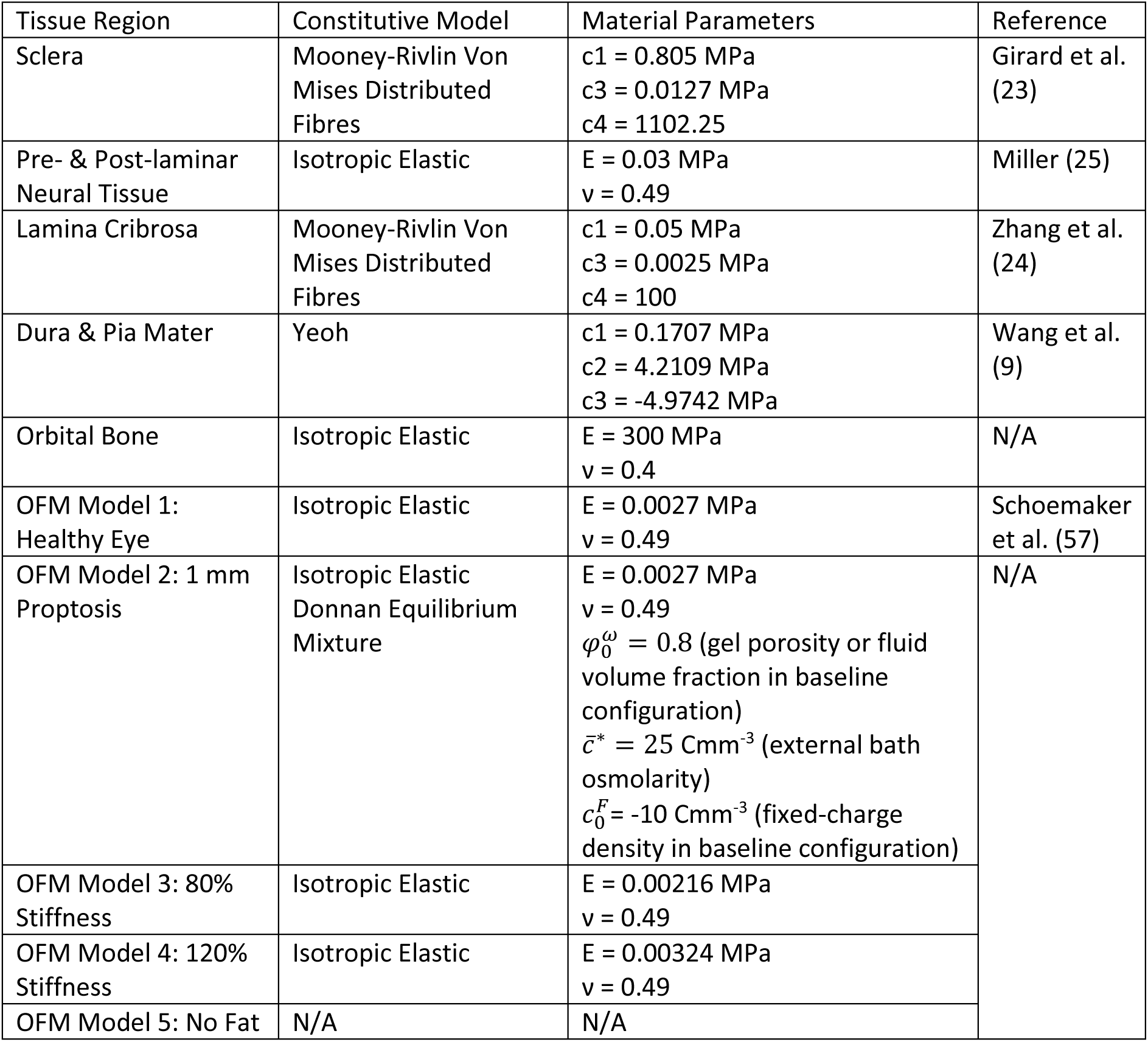
List of Material Properties.

| Tissue Region | Constitutive Model | Material Parameters | Reference |
| --- | --- | --- | --- |
| Sclera | Mooney-Rivlin Von Mises Distributed Fibres | $c1 = 0.805 \text{ MPa}$<br>$c3 = 0.0127 \text{ MPa}$<br>$c4 = 1102.25$ | Girard et al. (23) |
| Pre- & Post-laminar Neural Tissue | Isotropic Elastic | $E = 0.03 \text{ MPa}$<br>$\nu = 0.49$ | Miller (25) |
| Lamina Cribrosa | Mooney-Rivlin Von Mises Distributed Fibres | $c1 = 0.05 \text{ MPa}$<br>$c3 = 0.0025 \text{ MPa}$<br>$c4 = 100$ | Zhang et al. (24) |
| Dura & Pia Mater | Yeoh | $c1 = 0.1707 \text{ MPa}$<br>$c2 = 4.2109 \text{ MPa}$<br>$c3 = -4.9742 \text{ MPa}$ | Wang et al. (9) |
| Orbital Bone | Isotropic Elastic | $E = 300 \text{ MPa}$<br>$\nu = 0.4$ | N/A |
| OFM Model 1: Healthy Eye | Isotropic Elastic | $E = 0.0027 \text{ MPa}$<br>$\nu = 0.49$ | Schoemaker et al. (57) |
| OFM Model 2: 1 mm Proptosis | Isotropic Elastic<br>Donnan Equilibrium Mixture | $E = 0.0027 \text{ MPa}$<br>$\nu = 0.49$<br>$\phi_0^\omega = 0.8$ (gel porosity or fluid volume fraction in baseline configuration)<br>$\bar{c}^* = 25 \text{ Cmm}^{-3}$ (external bath osmolarity)<br>$c_0^F = -10 \text{ Cmm}^{-3}$ (fixed-charge density in baseline configuration) | N/A |
| OFM Model 3: 80% Stiffness | Isotropic Elastic | $E = 0.00216 \text{ MPa}$<br>$\nu = 0.49$ | |
| OFM Model 4: 120% Stiffness | Isotropic Elastic | $E = 0.00324 \text{ MPa}$<br>$\nu = 0.49$ | |
| OFM Model 5: No Fat | N/A | N/A |  |

### Boundary and Contact Conditions

To appropriately constrain the model, the outer surface of the orbital boundary was spatially fixed. The posterior terminus of the optic nerve was also fixed, to account for the adhesion of the optic nerve to the bones of the optic canal.

To generate an eye movement, two regions approximating the horizontal rectus muscle insertions on the corneo-scleral shell were rotated by an applied angular displacement of 13° about the centre of the eye globe, producing an effect analogous to a horizontal forced duction. As with our previous work, this rotation magnitude was chosen to match the eye movement recorded in our MRI data that was used to construct the FE model. Both adduction (nasal rotation) and abduction (temporal rotation) was simulated.

As in our previous model, a frictionless sliding interface was assumed between the posterior sclera and orbital fat to simulate the effects of Tenon’s capsule (26). Contact between the pia and dura was also assumed to be frictionless. The optic nerve and surrounding OFM were tied at their interface, based on prior studies (27) and our observation from dynamic MRI. The OFM was allowed to slide against the orbital boundary with a frictional coefficient of 0.5 (28).

An IOP of 15mmHg (applied to the inner limiting membrane) (29), a CSFP of 12.9 mmHg (applied within the subarachnoid space) (30), and an orbital tissue pressure of 4.4 mmHg (applied to the outer surface of globe and optic nerve) (31) were used as loading conditions in the FE model (**Figure 2**). These values represent averaged normal pressures in the supine position from which we obtained our MRI data.

**Fig 2:**
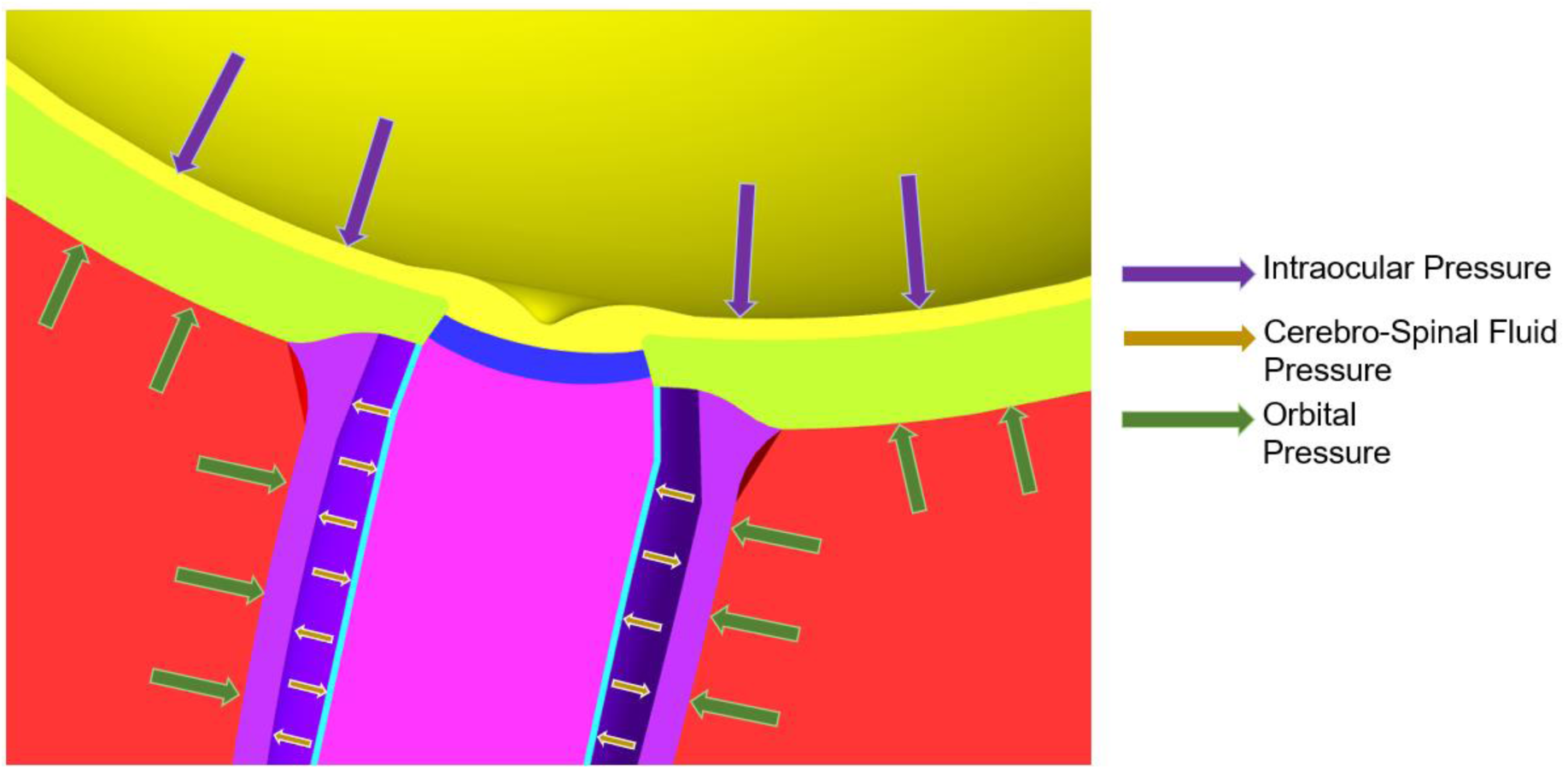
Illustration of the various pressure loads in all models.

### Differences between Eye Models in Healthy and TED Conditions

The above description of the eye models lists the properties that are common to all models. Here we clarify the differences between the five models, which comprise various modifications to the OFM material. (Table 1).

#### Model 1 – Eye Movements in a Healthy Eye

The OFM is characterized as an isotropic elastic material with a shear modulus of 900 Pa. This model is identical to one used in our earlier work (9) to simulate movements in a normal eye.

#### Model 2 – Eye Movements in an Eye with Orbital Tissue Swelling

In an estimated 60% of TED cases (32, 33), the volume of orbital tissue (either orbital fat or extraocular muscle) increases, causing exophthalmos. To model this, the OFM in the second eye model was characterized as a Donnan-equilibrium solid. In this material model, the solid is defined as a charged matrix immersed in a solution of countervalent ions. Altering the charge density of the solid matrix causes uptake of the surrounding fluid. This is not intended to be a perfect description of infiltrative orbitopathy on the micro-scale, rather, it is a mechanical approximation chosen to produce a swelling effect within the material. The effect of this material model applied to the OFM was that 1 mm of proptosis was induced (**Figure 3**). We expected some mechanical effects to be observable at this apparently minor displacement, as a difference in exophthalmometry measurement as small as 2 mm between eyes is considered to be diagnostically relevant for TED (34, 35).

**Fig 3:**
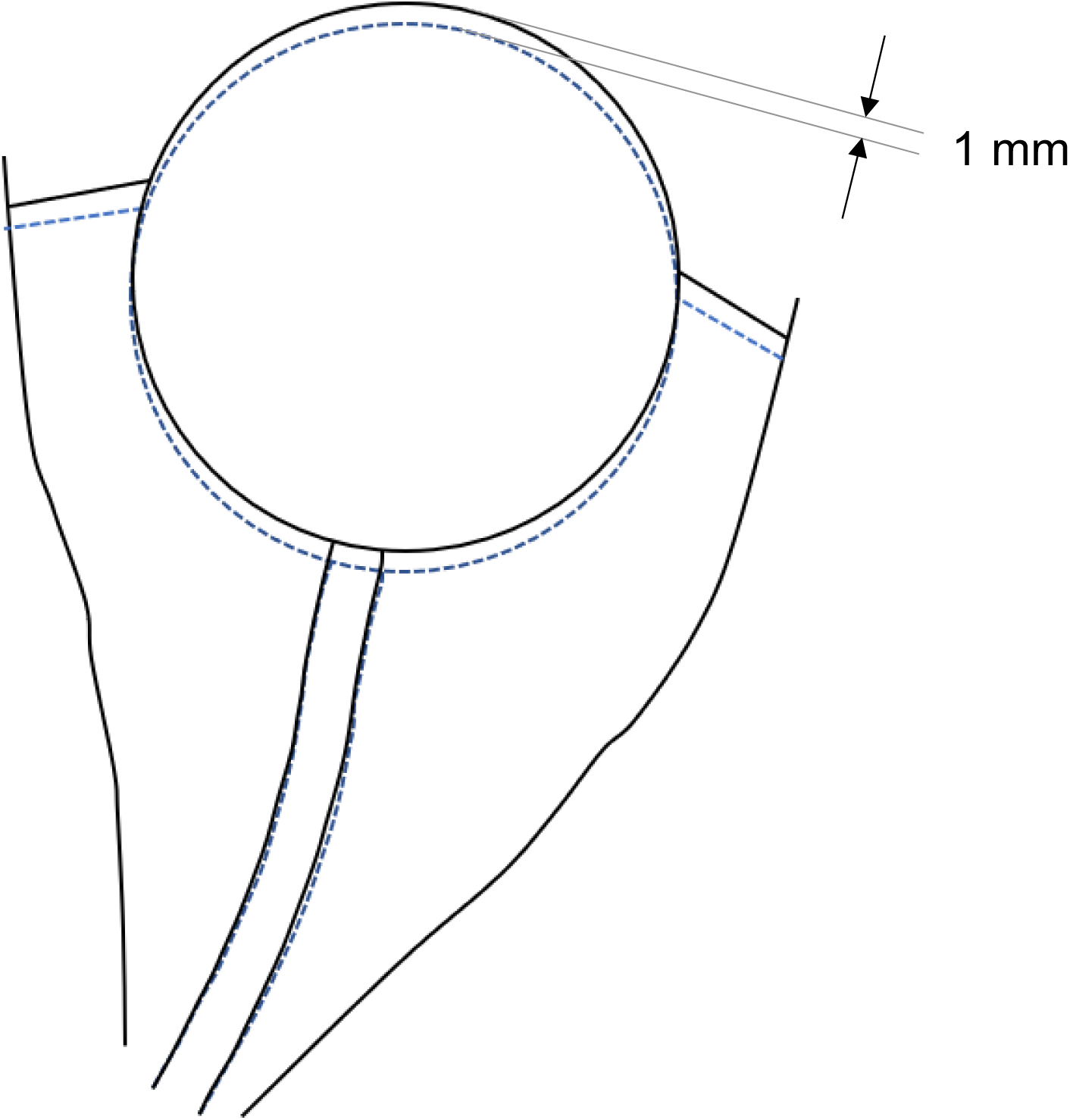
2D cross-section through the mid-transverse plane of the model orbit illustrating the geometry in baseline gaze. The dashed blue line illustrates the healthy geometry and the solid black line illustrates the geometry after swelling of the OFM material induces 1 mm of proptosis.

#### Model 3 and 4 - Eye Movements in an Eye with Orbital Tissue Stiffness Variation

In as many as 25% of TED cases there is no volume increase of orbital tissue, but the orbit may still be subject to fibrotic or infiltrative changes (36). To reflect this, in the third model the OFM was characterized as an isotropic elastic material, similar to the first (normal) model, but the shear modulus was increased by 20%. A fourth model was similarly constructed with a 20% decrease in stiffness. This fourth model was not intended to reflect any specific disease state but was included to better illustrate the effects of variation in the OFM.

#### Model 5 – Eye Movements in an Eye lacking Orbital Soft Tissue

The final eye model had all orbital soft tissue removed, such that the globe and optic nerve appeared unsupported in empty space. This apparent physiological impossibility is equivalent to assuming that the orbital fat environment does not affect the motion of the nerve. Some previous models have omitted the orbital fat when performing mechanical simulations at the ONH (11, 37), and it may be important to understand the implications of this assumption.

### Mechanical Analysis of the Numerical Eye Models

To assess the effects of TED on the mechanics of the ONH, measures of deformation are required. We used a measurement known as the tilt angle, which has recently been introduced to measure ONH deformation in OCT images (15). The tilt angle measures relative antero-posterior displacement of the nasal and temporal sides of the Bruch’s membrane opening (BMO) in eye movement. For a detailed description of how this measurement is performed, refer to the supplementary material.

Additionally, for each FE model we reported the effective stress in the peripapillary sclera), reflecting the local average internal force experienced by this region. Separate stress measurements were made for the nasal, temporal, and superior/inferior quadrants (**Figure 5**).

**Fig 4:**
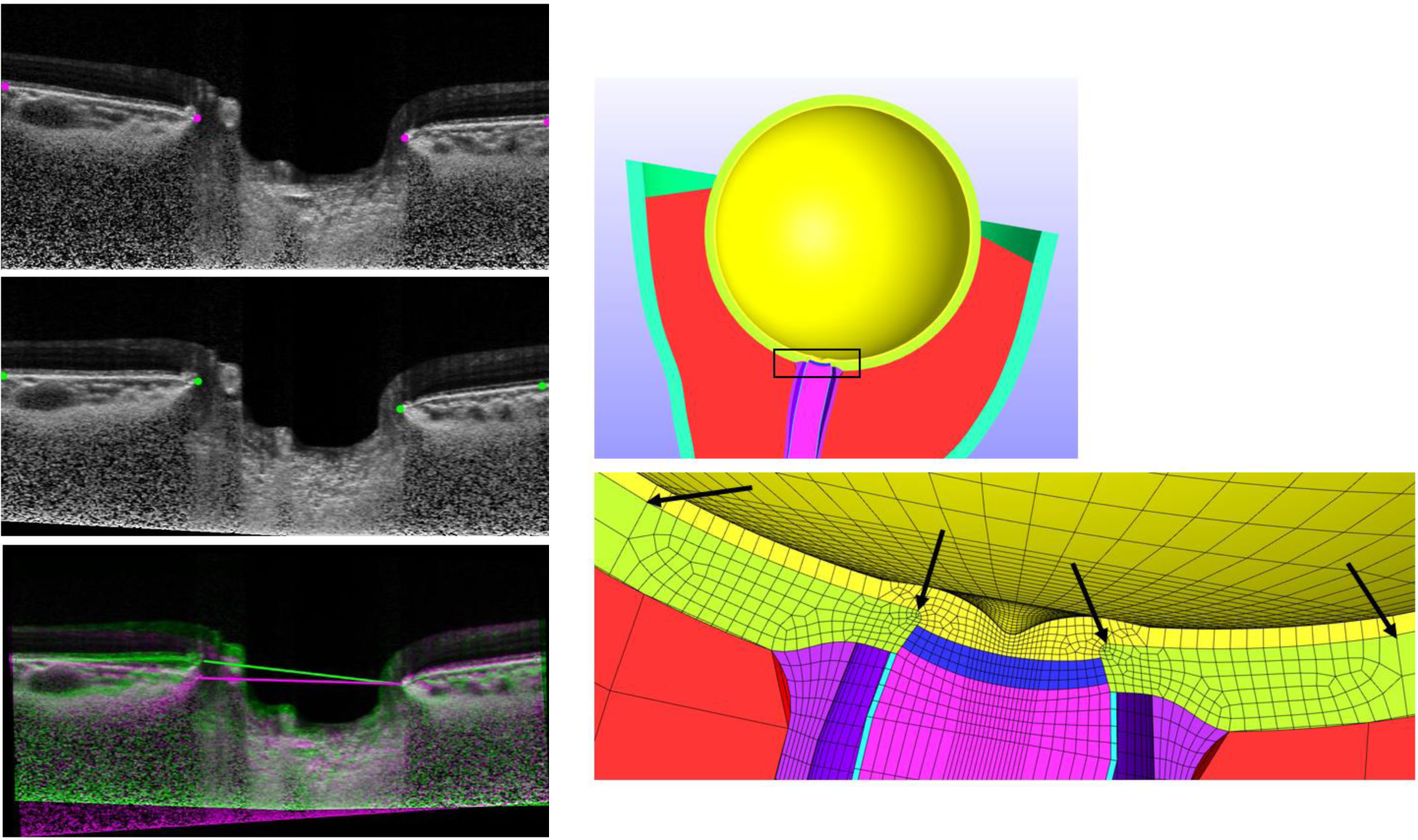
Procedure for tilt angle measurement. Central OCT B-scans in primary gaze (top left) and 20 degrees of eye movement (middle left) are marked for peripheral BM points and BMO points. Images are rotated to align these peripheral BM points (bottom left) and then the tilt angle can be measured as the angle between the lines connecting the BMO in each image. This procedure was adapted for numerical simulations by recording the displacement of corresponding locations in the finite-element mesh (bottom right, arrows). For the in vivo measurements these points must be manually marked on the OCT images. In the finite element models the location of the mesh points is computed directly by the numerical solver.

**Fig 5:**
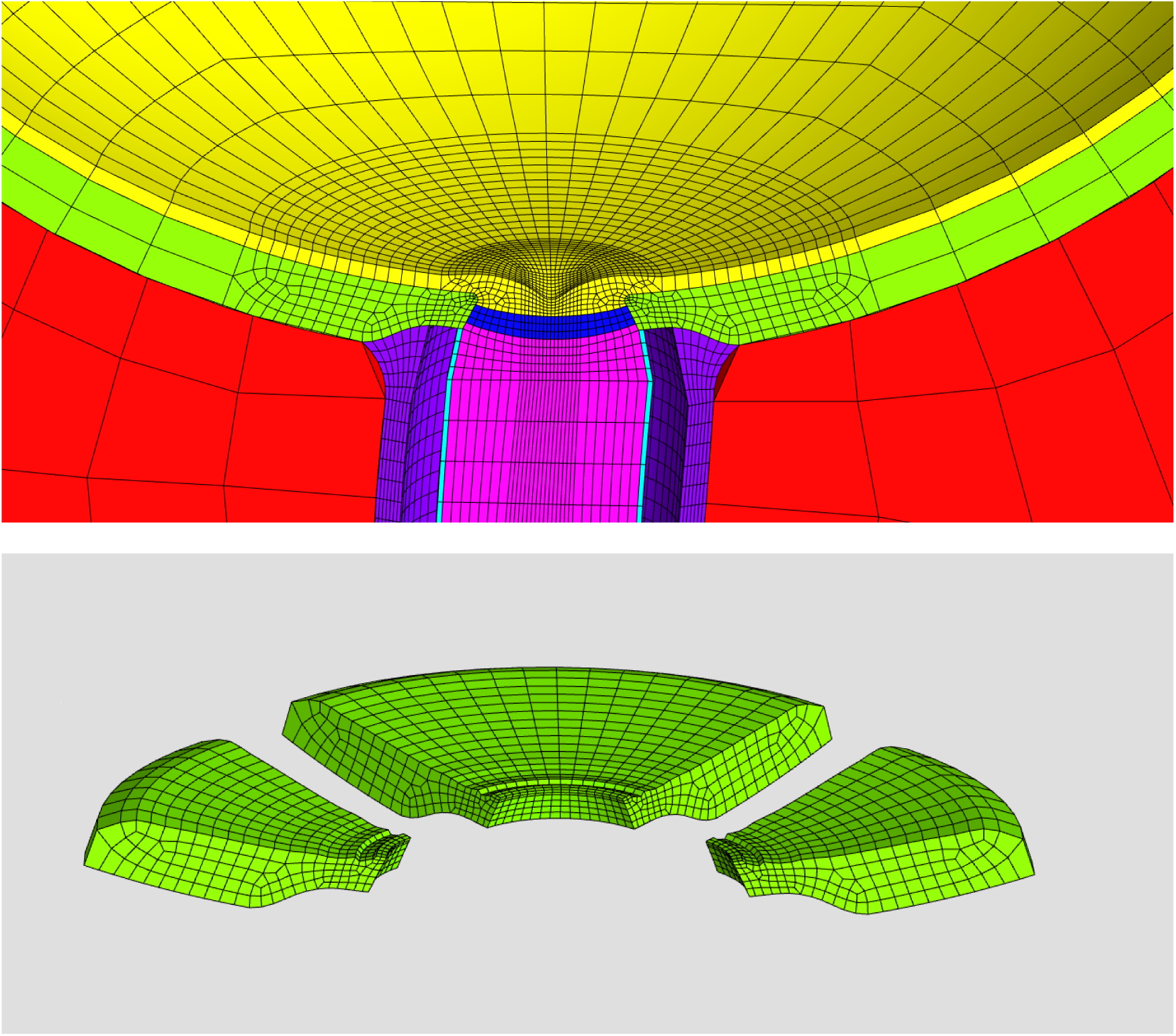
(Top:) The optic nerve head region of the finite-element model. Visible structures are the prelaminar nerve tissue (yellow), sclera (green), lamina cribrosa (blue), postlaminar nerve tissue (magenta), pia mater (cyan), dura mater (violet). and orbital fat (red). (Bottom:) Exploded view of the peripapillary sclera illustrating nasal, superior/inferior, and temporal (left-to-right) regions. An anterior layer of the sclera elements has been removed in this illustration: these elements were ignored in the stress analysis to highlight the posterior effects of the orbital fat and optic nerve on the sclera.

### OCT Imaging in Baseline Gaze Position

To confirm our computational results with clinical observations, one patient with TED and 10 healthy subjects were recruited from the Singapore National Eye Centre. All subjects gave written informed consent. The study adhered to the tenets of the Declaration of Helsinki and was approved by the institutional review board of the Singapore National Eye Centre.

The TED patient had bilateral severe disease with optic neuropathy manifested by reduced visual acuity, visual field deficit, and restricted extraocular movements. IOP in both eyes was normal (with assistance of prescribed IOP-lowering eyedrops). Images were only recorded in the patient’s right eye due to the constraints of the clinical protocol.

The measurements of the cohort of healthy patients were performed for a prior study (13), and the data were analysed retroactively. For the healthy cohort, OCT measurements were recorded in both eyes.

To compare with the images of the ONH in eye movement, an initial OCT volume of each subject’s ONH was obtained in the baseline OCT position, using spectral domain OCT (Spectralis; Heidelberg Engineering GmbH, Heidelberg, Germany). Note that for all subjects, this imaging of the ONH was not strictly performed in primary gaze, as the relative position of the subject’s head and the OCT target in the ‘default’ orientation requires the subject to perform a small adduction movement of approximately 7 degrees. Therefore, in this paper, we use ‘baseline gaze position’ to refer to this slightly adducted eye position during a standard OCT scan. The in vivo eye rotations reported in this study were measured relative to this baseline gaze position.

### OCT Imaging Following Changes in Gaze Position

Two additional OCT volumes for each subject were obtained in different gaze positions: one with the eye in nasal gaze and one with the eye in temporal gaze. Both rotations were 20° from the baseline position. Eye rotation was achieved by rotating the subject’s head while keeping the eye aligned with the stationary OCT objective. Subjects were positioned on a custom 3D-printed chin rest for precise control of the head rotation.

### Mechanical Analysis of In Vivo Images

To assess the amount of deformation occurring at the ONH in eye movement in vivo, we used the same ‘tilt angle’ measurement technique described previously. (Refer to supplementary material). (Figure 4).

## Results

### Measurements of tilt angle in model eyes

In the FE model of the normal eye, tilt angles were 6.26° in adduction and −5.91° in abduction. These angles increased (6.45° in adduction and −6.18° in abduction) when the stiffness of the OFM was raised by 20% and decreased (5.93° in adduction and −5.55° in abduction) when this stiffness was reduced by 20%. In the eye model with increased orbital tissue volume and proptosis, the tilt angles were greater (10.02° in adduction and −10.90° in abduction). In the model lacking OFM material the tilt angles were smaller (4.33° in adduction and −2.93° in abduction). Angles in abduction were always negative (the nasal side of the BMO moved posteriorly relative to the temporal side) and the angles in adduction were always positive. (**Figure 6**).

**Fig 6:**
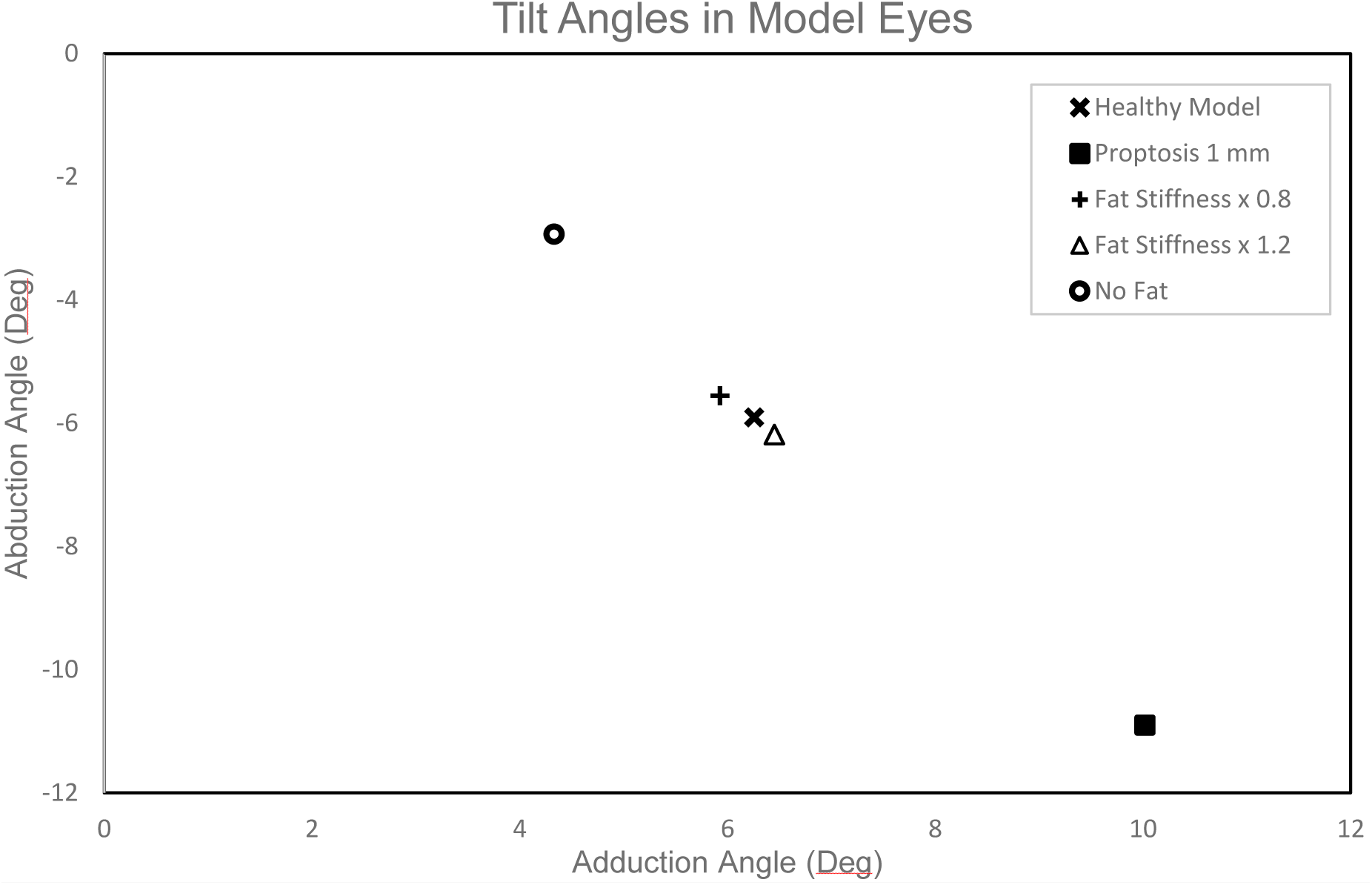
Tilt angles measured from finite element simulation. A negative angle indicates that the nasal side of the BMO moved posteriorly relative to the temporal side, and vice versa.

### Measurements of peripapillary scleral stress in model eyes

In all models, the maximum effective stress was observed in the quadrant contralateral to the eye movement direction (i.e. for adduction, or nasal rotation, the maximum stress was in the temporal quadrant, and vice versa). Stress in the superior/inferior quadrants was an intermediate value and stress in the ipsilateral quadrant was lowest. (**Figure 7**).

**Fig 7:**
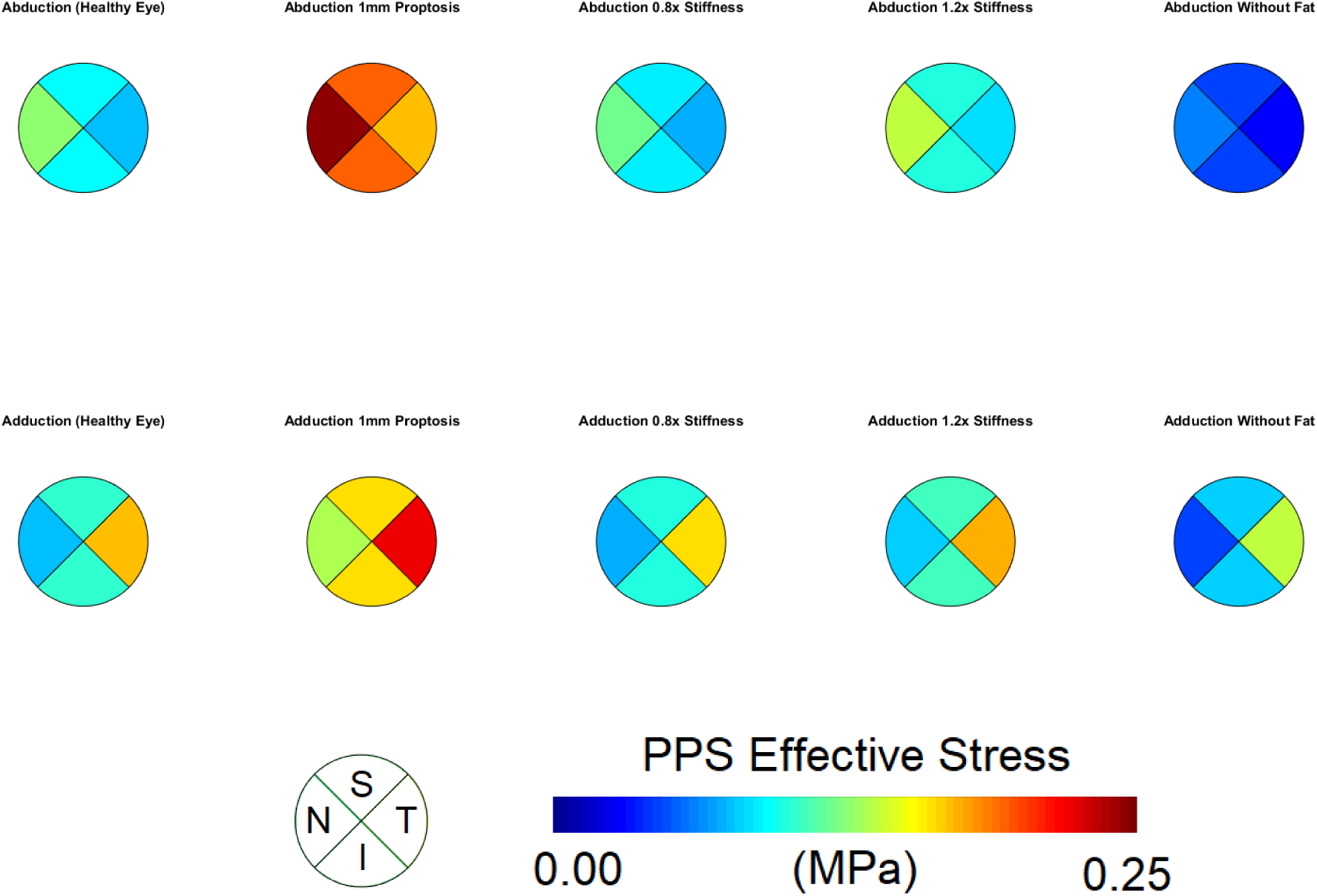
Colour map of effective stress in the peripapillary sclera of the model eyes following eye movement. Measurements are averaged over the nasal (N), temporal (T), superior (S) and inferior (I) quadrants. (See figure 5). Due to the model symmetry the measurements in the superior and inferior quadrants are equal.

The model with the lowest stress was the 5^th^ model with no orbital tissue. The first, third, and fourth models (the normal eye and the two eyes with OFM stiffness adjusted by 20%) had intermediate stress values. In these three models the stress increased slightly with greater tissue stiffness. The models with swollen orbital tissue exhibited the greatest stress. (**Figure 7,8**)

**Fig 8:**
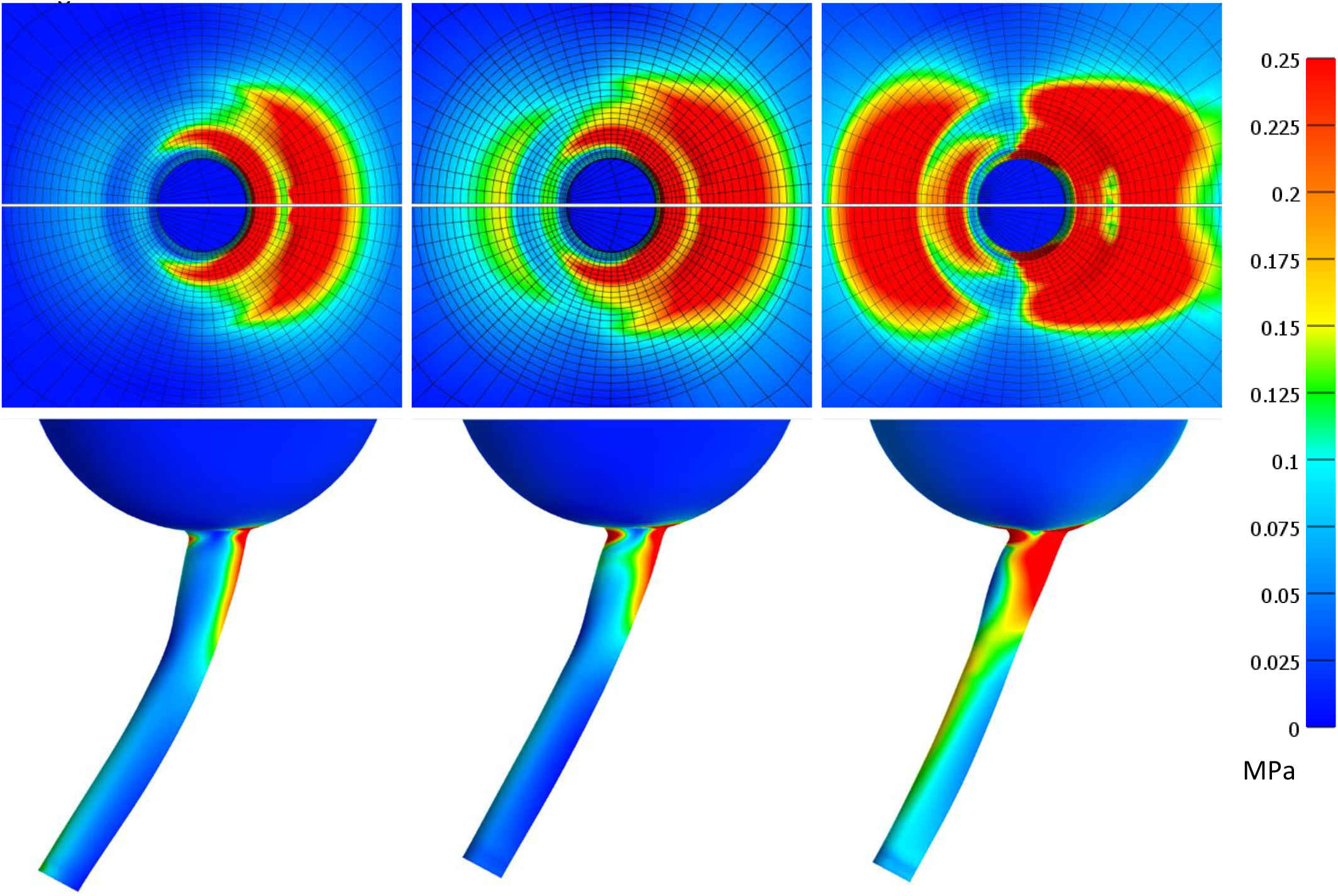
Colour map of effective stress in the posterior sclera and optic nerve sheath following eye movement (adduction). Left-to-right images show model 5 (No fat), model 1 (normal eye) and model 2 (1 mm proptosis).

In general, stress measurements were greater in adduction than for the corresponding model in abduction. The exception to this was the model with the swollen orbital tissue, for which stresses in abduction were greater than adduction. This was also the only model eye with a greater tilt angle in abduction than adduction.

### In vivo measurements of tilt angle in human eyes

For the human eyes, tilt angles measured in adduction were also always positive, meaning that the temporal BMO moved posteriorly relative to the nasal BMO. The tilt angle for healthy eyes in adduction was 1.46° ± 0.25 (mean ± standard error of the mean). In most eyes the tilt angles in abduction were negative (i.e. the nasal BMO moved posteriorly relative to the temporal BMO), but in some eyes there were positive tilt angles in both abduction and adduction. The mean tilt angle for healthy eyes in abduction was −0.42° ± 0.12. For each eye, the magnitude (absolute value) of tilt angle in abduction was typically lesser than in adduction, with the Wilcoxon signed rank test rejecting a median difference of zero (p < 0.001). A linear correlation was observed between the magnitudes of tilt angle in each movement direction in the healthy eyes, as eyes with larger positive tilt angles in adduction tended to also have larger negative tilt angles in abduction (R^2^ = 0.24). The eye with thyroid eye disease had the largest tilt angles in both adduction and abduction (5.37° and −2.21° respectively). (**Figure 9**)

**Fig 9:**
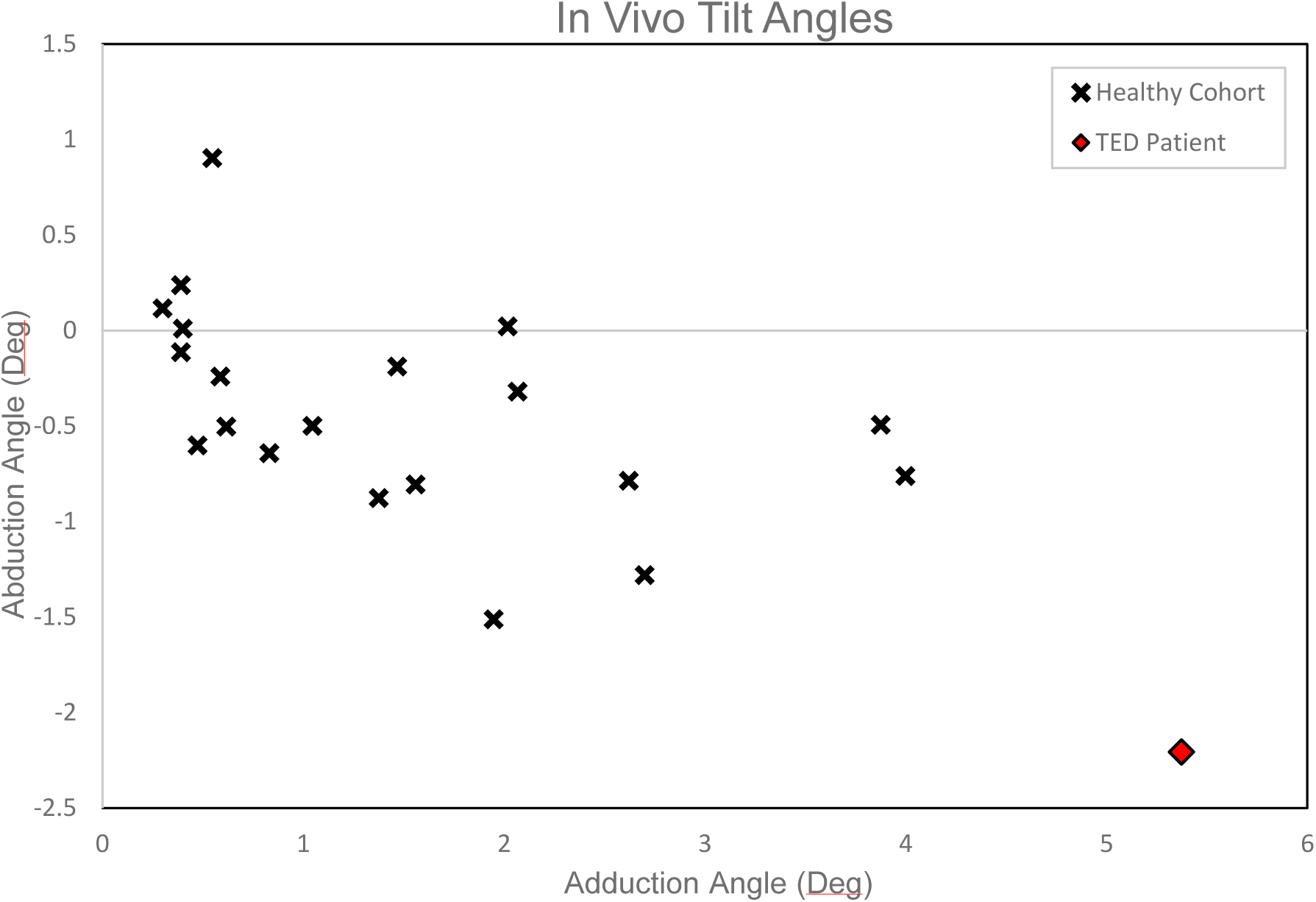
Tilt angles measured in vivo. Each marker represents one eye. A negative angle indicates that the nasal side of the BMO moved posteriorly relative to the temporal side, and vice versa.

## Discussion

Contemporary research suggests that gaze-evoked deformations caused by elevated optic nerve tractions may lead to glaucoma-like injury. TED may worsen these tractions due to proptosis or exaggerate the deformations by altering the mechanical properties of the surrounding orbital fat. Numerical results from FE models support this theory and it is further strengthened with our gaze-evoked deformation measurements in our TED patient. This suggests that orbital fat is an important structure to consider in the context of eye movements or ocular biomechanics generally.

### Comparison of Tilt Angles

The FE models showed a slight increase in tilt angle when the elastic modulus (stiffness) of the OFM tissue was increased by 20%. In contrast, the increase in tilt angle when 1mm of proptosis was induced was greater. This may seem to imply that the stiffness of the orbital fat has a limited influence on the ONH compared to the amount of proptosis. However, in model 5 where the orbital tissue was removed entirely, the tilt angle measurements decreased greatly, even though the exophthalmometry was the same as the healthy model. To further explore this relationship, we created an additional model without OFM tissue, and induced 1 mm of proptosis by directly manipulating the position of the globe. Tilt angle measurements in this model were slightly greater than the original no-OFM model (4.65° in adduction and −2.75° in abduction), but still lesser than the tilt angles in the healthy model with normal exophthalmometry and OFM material. Additionally, the swelling of the Donnan material model involves an elastic deformation of the underlying solid matrix, which will influence its response to subsequent applied loads. Essentially, the swelling of the Donnan material is a combined volume increase and stiffening process. If proptosis alone does not cause a great increase in tilt angle (as was the case comparing the models with no OFM material) then the stiffness of the orbital fat explains the large difference in tilt angles between models 2 (swelling OFM) and 5 (no OFM). The reason why this large difference was not observed in models 3 and 4 is likely because adjusting the elastic modulus by 20% had a much lesser impact on the effective stiffness of the OFM than applying the swelling or removing the material entirely.

While both the models and the human measurements showed that tilt angles could be increased in TED, the tilt angles recorded in our human cohort were smaller than those in the FE model. The greatest tilt angles in the human population (the TED eye) were comparable to the smallest angles in any model eye (the model without OFM material), even though the simulated eye movement in the model eyes was smaller than the movement in the human eyes (13° vs 20°). This finding suggests that our model of a normal eye could be improved. It is not known which aspect of the model (material properties, geometry, boundary conditions, etc) is causing this discrepancy, as there are multiple ways that the tilt angle in the model could be reduced (e.g. reducing the OFM stiffness or increasing the scleral stiffness would both have this effect).

Sibony has measured much greater in vivo tilt angles of approximately 8-15° (in 30° eye movements), but these data were from a population of eyes with papilledema so the results are not directly comparable.(22)

### Optic nerve traction as a basis for vision loss

At present the link between gaze-evoked deformations and the development of chronic ocular pathologies is largely speculative. It has been argued that the deformation of ocular tissues in eye movements are more severe than the deformations caused by glaucoma-inducing IOP elevations (9). The prevailing explanation for these deformations is that the optic nerve applies a traction to the posterior pole of the globe when the ONH moves away from the orbital apex in eye movement. This geometric explanation was suggested by Friedman as early as 1941 (38). However, it is not known whether ONH deformation in eye movement and in glaucoma differs qualitatively in some important aspect, or whether the transient deformations of eye movement should have similar long-term effects as the constant forces of high IOP. New research has observed a higher incidence of globe retraction during horizontal eye movements in some cases of glaucoma (39). It is possible that this indicates that these patients are experiencing greater optic nerve tractions, and therefore more significant gaze-evoked deformations, contributing to their loss of vision. Repeated saccadic motions could have damaging effects analogous to overuse syndromes.

On the other hand, the link between greater optic nerve tractions and neurological damage is not widely accepted in TED. Some publications do support the idea of ‘stretch neuropathy’, where proptosis in TED damages the nerve by axial stretch (40-43), but conflicting evidence suggests that dysthyroid optic neuropathy (DON) has no connection to larger exophthalmometry measurements, shorter optic nerve length, or lesser optic nerve tortuosity (44). These factors should exacerbate optic nerve tractions, and therefore under the hypothesis of harmful gaze-evoked deformations they should be associated with neuropathic outcomes. The standard explanation for the etiology of DON is instead compression of the optic nerve at the orbital apex by expansion of the extraocular muscles. It is well-established that optic nerve traction does cause vision loss in acute proptosis, such as in trauma cases (45-47). However, sceptics of the ‘stretch neuropathy’ hypothesis in TED have proposed that the optic nerve can safely accommodate increased traction when inflammation causes proptosis to develop gradually (46, 48).

These two apparently contradictory positions can be reconciled by speculating that tractions in TED do not cause the same neuropathy as apex muscle crowding, but instead lead to an eventual glaucoma-like injury. Evidence of this occurring in TED might be a pattern of visual field loss that is atypical for DON or similar to glaucoma, or visual loss that cannot be reversed by decompressive procedures. The pattern of visual field loss in compressive thyroid neuropathy has recently been described as distinct from that in glaucoma (49). There have also been isolated reports of ‘atypical’ field losses in TED (such as inferior altitudinal defects (50) or generalised constriction (51)) that could not be successfully addressed by decompression, but there is disagreement as to how unusual these field losses are, with other studies suggesting that inferior altitudinal defects are a sign of late-stage DON (where visual losses might be expected to be irreversible) (49) or that field constriction in TED is relatively common (52).

If we hypothesize that nerve traction in TED leads to a glaucoma-like injury then we would expect to find an association between TED and glaucoma, but this relationship is complicated by the possibility of TED causing ‘true’ glaucoma by elevating IOP. It has been variously reported that TED is associated with high IOP but not glaucoma (8), that TED is associated with high IOP and that glaucoma is associated with orbitopathy duration (53), that TED is associated with glaucomatous RNFL changes (54), and that TED is associated with high IOP, open-angle glaucoma, and normal-tension glaucoma (7). This latter finding that TED is associated with normal-tension glaucoma invites optic nerve traction as a possible cause in these cases, but the studies have found no relation between glaucoma and proptosis measurement (7, 53), again suggesting that nerve traction is perhaps not a contributing factor.

### Non-traction explanation for gaze-evoked deformations

The leading explanation for gaze-evoked ONH deformations is optic nerve tractions: when the globe has sufficiently rotated such that the slack in the nerve is eliminated, then any further rotation will apply a pulling force to the optic nerve head. It has been determined that an angular threshold for this pulling effect is approximately 26° in normal subjects in adduction (15), but some deformation occurs during rotations that are lesser than this, so an additional explanation is required.

One suggestion is that the inherent stiffness of the optic nerve or ON sheath resists bending, so that even if the nerve is not completely taut it still applies a force to the ONH whenever the eye rotates. Numerical simulation has predicted that stiffer dura and pia mater increases lamina cribrosa strains during eye movement (9).

Another possibility we suggest here is that the presence of orbital fat obstructs the motion of the optic nerve, and that a resultant shear or torque is applied to the ONH during eye movements, exacerbating gaze-evoked deformations. (**Figure 10**). Our simulations indicate that both the presence of the orbital fat and its stiffness affect gaze-evoked deformations. Orbital fat in TED has been previously noted to have elevated stiffness (55), and our measurements of a TED eye indicated greater gaze-evoked deformations. This fat obstruction is a plausible explanation for why the TED patient in this study had greater gaze-evoked ONH deformations than the healthy cohort. Elevated optic nerve tractions may not be a sufficient explanation for this finding, because the exophthalmometry measurements for this patient were within normal limits (34, 35, 56).

**Fig 10:**
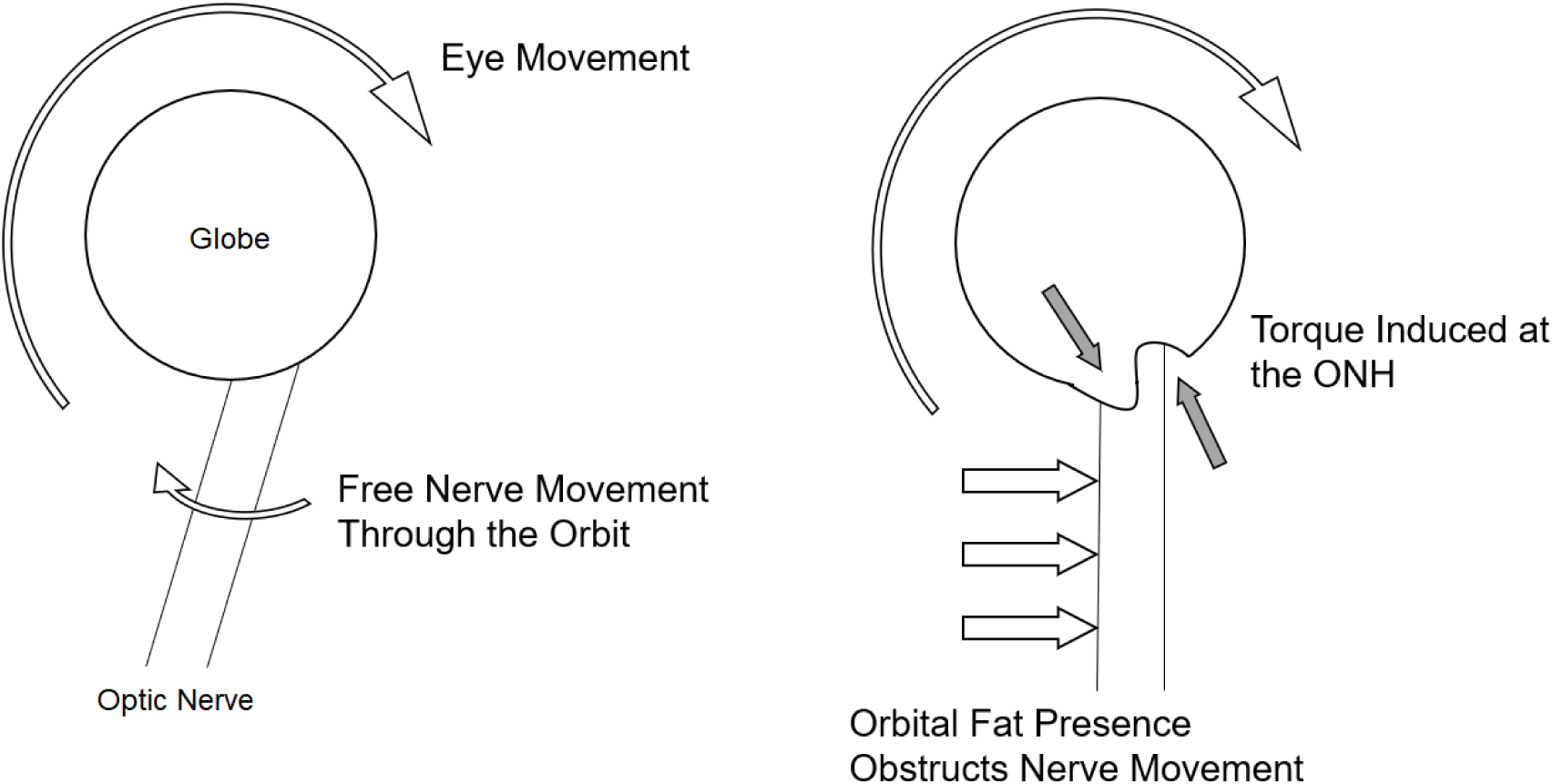
(Left) An eye movement in an ‘empty’ orbit. As the eye rotates, the optic nerve is free to traverse the orbit without resistance. (Right) When the orbit contains elastic fat, motion of the nerve in eye movement may be impeded by the fat. The optic nerve then would act as a lever arm, delivering this resistance to the ONH as a torque that causes the ONH to tilt. This effect may be magnified in TED when the stiffness of the orbital fat increases.

These observations are complicated by the fact that we have no direct measurement of the orbital fat properties of this TED case, and that other independent mechanical factors (such as orbital congestion by muscle involvement) could have also affected gaze-evoked deformations in this case. However, this preliminary evidence suggests that gaze-evoked deformations in TED may deserve further investigation.

## Limitations

Our TED patient did have noticeably larger tilt angles than a sample from the normal population, and we suspect this is attributable to the orbital disease, but there are many ways that TED can present, and these various manifestations may affect gaze-evoked deformations independently. A larger sample of TED patients at different disease stages would lead to a more complete understanding of this finding.

The model simplification of intraorbital tissue as an ‘OFM complex’ lacking discrete muscle structures was considered an appropriate measure, due to the region of interest being the immediate ONH region located some distance from the muscle bellies in a real orbit. However, in TED the muscle and fat can be differentially involved, so it may be interesting to study gaze-evoked deformations when the properties of fat and muscle are altered separately. The creation of such a model is a future objective.

Similarly, our Donnan model of inflamed tissue is just one of multiple potentially valid numerical choices for describing TED, as the various possible orbital manifestations in TED mean that a specific proptosis measurement does not represent a unique mechanical state. Many combinations of globe position and material properties could potentially be simulated, but we limit ourselves to these few general models to introduce some novel mechanical consequences of TED.

## Supplementary Material: Measuring ONH Deformation In Vivo

In the field of ocular biomechanics there has been recent interest in measuring the in vivo deformation of ocular tissues during eye movement. It is established that ONH tissues can be visualised using OCT but quantifying the displacement of these tissues between successive images is challenging. The entire field of an OCT image of the posterior segment contains easily deformable soft tissue, and capturing an image of an eye in an adducted or abducted position requires the subject to perform a large movement of their head to realign their eye with the OCT fixation target. These factors introduce image registration difficulties.

The deformation measurement used in this paper is the tilt angle, which measures relative anteroposterior displacement of the nasal and temporal sides of the Bruch’s membrane opening. To perform this measurement in the FE models, mesh nodes at the nasal and temporal points of the BMO are noted (the points where the Bruch’s membrane ends at the opening of the scleral canal) in the horizontal cross-section through the centre of the ONH. A line can be constructed through these two nodes in the baseline gaze position. After eye movement, the new position of the globe can be registered to the initial position and a second line can be constructed between the same two nodes. The tilt angle is recorded as the angle between these two lines. (Both lines always lie in the transverse plane, as we limit the analysis to horizontal eye rotations).

The choice of registration method is crucial, as the tilt angle measurement should only capture local deformation of the ONH tissues, but the absolute position of these BMO points in space is also affected by the bulk rotation of the eye globe. To control for this rotation, it is necessary to make the tilt angle measurement with reference to a point that does not deform during eye movement. It has been previously assumed (1) that the ocular tissues far from the ONH (close to the ocular equator) deform minimally during eye movement, as the likely source of gaze-evoked deformation is the force applied by the nerve at the ONH. Registration has therefore previously been performed in vivo by noting that for an OCT image centred on the ONH, the tissues at the extreme margins of the image are furthest from the ONH. The tilt-angle can then be measured by aligning the baseline and movement images at the two points where the BM intersects the lateral image boundaries.

For our FEM model, there was no limited image frame to constrict the field of view around the ONH, so we could have measured the tilt-angle deformation relative to some distant, fixed anatomical feature, such as the orbital bone. However, we intended to verify our model findings using a similar in vivo technique, so we selected nodes in the mesh at the approximate location of where the BM would intersect the image frame in our OCT images. The tilt angles were measured after rotating and translating the mesh coordinates so that the position of these nodes coincided, thereby reproducing the in vivo method (**Figure 4**, **main manuscript**).

For the OCT of the human subjects we have used a similar technique, also assuming that there was minimal deformation in tissue far from the ONH, the same assumption also used by Suh et al. to measure the BMO tilt angle by registering images at the ‘intersection points’ where the BM intersects the image boundary (1). While we agree that this assumption is reasonable, we observed that in several cases the image boundary did not intersect the Bruch’s membrane at similar locations in successive images, presumably due to misalignment of the subject’s eye globe with the OCT system during eye movement. If registration is then performed at these ‘roving’ intersection points, artificial displacements are measured. To avoid this problem, we cropped the field-of-view of the OCT images so that the linear distance between these lateral endpoints and the respective BMO endpoint was the same in each image, effectively ensuring that the ONH was centred in each OCT image (**Figure S1**). While this step assumes that deformation of the peripapillary region is mostly tilt and does not include gross stretch or bending, inspection suggested that this step produced more accurate registration than always assuming the subject’s eye was always precisely aligned with the OCT device.

Other registration techniques have been reported in the literature; an earlier publication (2) also made the assumption that the intersection between the BM and image boundary deforms minimally, but rather than using intersection points on both the nasal and temporal sides of the image, they instead constrained image rotation by co-localising an intersection point on only one side of the image, then rotating the images so that the paths of the BM through this point are tangential. In this method, a change in curvature of one side of the BM will affect the apparent position of the BM on the opposite side, so the ‘tilt angle’ measurements are dependent on the side of the image chosen for registration. We agree with the later suggestion by these authors that this method is obsolete (1).

Lee et al. similarly assessed ocular deformation in eye movements using OCT, using the foveal dimple and the centre of the ONH as registration landmarks (3). Using the ONH centre (the midpoint of the BMO) as a registration point may introduce error, as this area is expected to be a region of significant deformation. However, the foveal dimple is a useful choice, as it is a clearly identifiable anatomical point far from the ONH. (Unlike the intersection between an image boundary and the BM, which does not represent a unique anatomical location). The limitation of this technique is that locating both the ONH and the foveal dimple in a single image requires the use of OCT that captures a wide view of the posterior eye. Furthermore, this method still relies on the same general assumption as the process we have used, as there is still no guarantee that these landmarks are stationary. Sibony has also made ‘tilt angle’ measurements but the method of image registration was not specified (4).

We also tested another angular-measurement technique whereby the points of the BMO themselves are used as fixed registration landmarks, such that the BM appears to pivot about the BMO in eye movement while the BMO remains stationary. Because images are registered to distinct points in the centre of the image field, this technique initially avoids the assumption that tissues deform minimally nearer the ocular equator. However, to measure the extent of this angular ‘BM displacement’ it is necessary to construct a ray from the BMO point to some endpoint on the BM. The position of this endpoint is either an arbitrary distance from the BMO, or at the intersection between the BM and the image boundary, which introduces similar problems as the other registration techniques. Also, the points of the BMO do move during eye movement, so this technique produces indirect measurements and does not describe the deformation in an intuitive way. On the balance of these considerations it was decided that this provided no meaningful advantage over the original ‘tilt-angle’ measurement.

More complex approaches for measuring deformation in tissues involve a quantification throughout the entire region of interest, rather than using discrete measurements such as tilt angle or cup depth. Prior examples for measuring deformation in eye movements include the use of geometric morphometrics (5, 6) and strain mapping (7). While these methods do provide a more detailed description of the deformation, registration is still similarly challenging, and it is still not known which (if any) aspect of transient ONH deformation contributes to negative physiological outcomes. It is assumed that large deformations from a primary-gaze baseline may have a detrimental effect, and it has previously been observed that the ‘seesaw tilt’ (relative anteroposterior displacement of the nasal and temporal peripapillary regions) is the most salient feature of deformation in horizontal eye movements (5). It follows that measurement of the tilt-angle may be a good approximation of overall deformation, and that conditions producing large tilt angles may be of clinical interest. We therefore only report measurements of tilt-angle, but a more detailed investigation of e.g. local strains may be warranted when there is greater understanding of how these tissues can be damaged by mechanical loading at the micro-scale.

**Fig S1:**
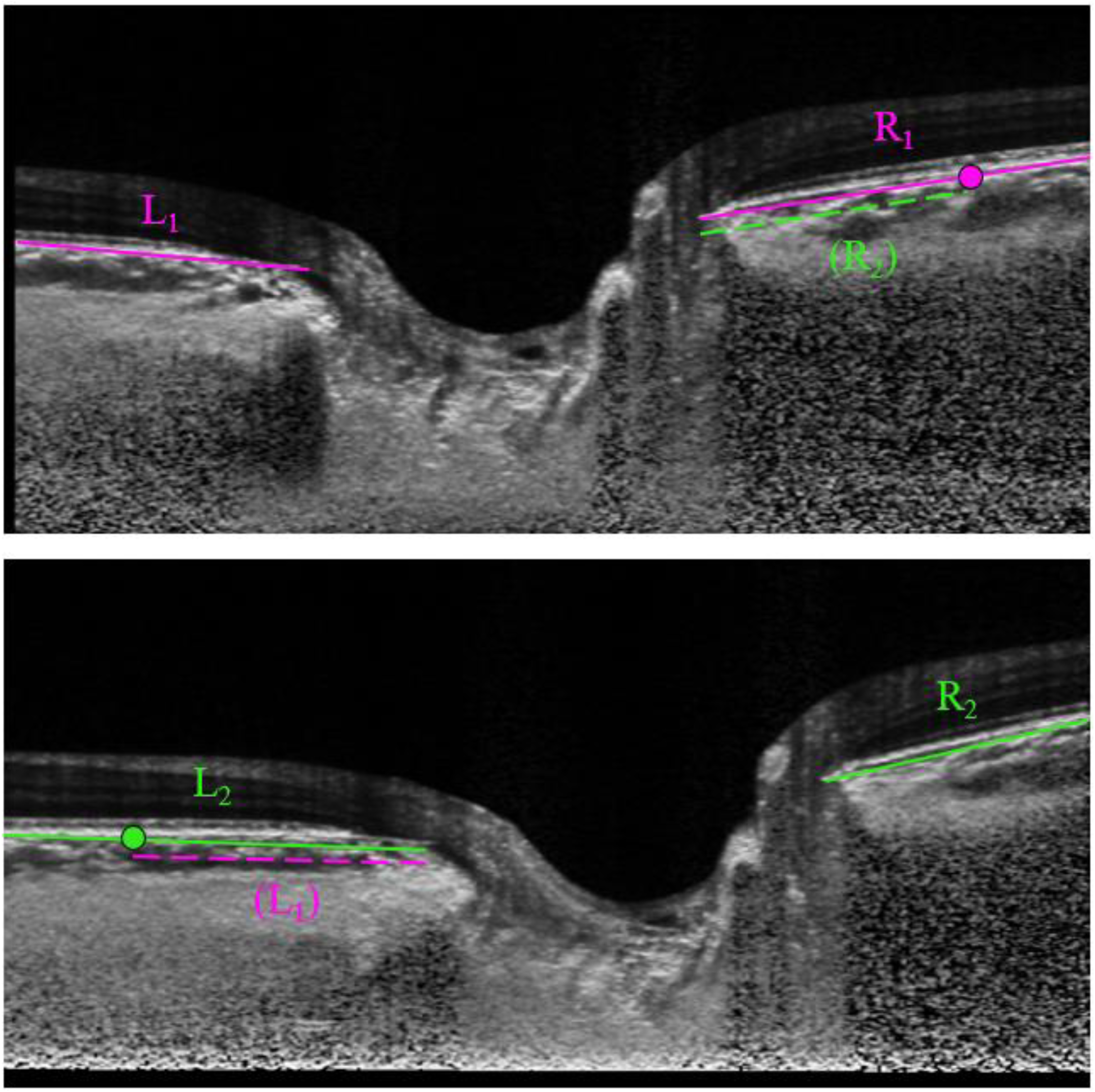
Two images illustrate baseline gaze (top) and eye movement (bottom). The optic cup appears to have shifted to the right in eye movement, but for this shift to be caused by local deformation the section of tissue visible to the left of the scleral canal needs to have stretched to approximately 1.5 times its original width. A more likely explanation for this observation is that the image frame is not similarly centred on the ONH in each image. Without any correction, the original registration method for the tilt-angle measurement (using points where the BM intersects the image frame) will lead to errors, as this inaccurate centring causes the image frame to intersect the BM at different anatomical locations. To correct for this, the linear distance from the BMO to the image frame is recorded for all four BM lengths (L1, L2, R1, and R2). The shorter length measurements of the two pairs (L1 and R2 in this example) are used to re-select registration points, effectively cropping the images so that they are both centred at the same position.

## Footnotes

1 ‘Adductions’ and ‘abductions’ are used in this paper to describe the rotation of an individual eye in the nasal or temporal direction, without considering any coordinated movement of the fellow eye. The mechanical effects would be identical in horizontal vergence or version movements of equivalent angular displacement.

